# Magnetic Bead Based Proximity Extension Assay for Sensitive Protein and Extracellular Vesicles Detection

**DOI:** 10.1101/2022.11.27.518072

**Authors:** Phathutshedzo Muthelo, Tonge Ebai, Ehsan Manouchehri Doulabi, Rasel A. Al-Amin, Yajun Wu, Masood Kamali-Moghaddam, Ulf Landegren

## Abstract

During the early stages of disease development, protein biomarkers can leak into the blood creating opportunities for early diagnosis of disease with minimally invasive sampling. These proteins biomarkers however are often masked by the presence of more abundant functional blood proteins, making specific detection a challenge with most current immunoassays. We we report on the development of a magnetic bead based solid-phase PEA (SP-PEA) for sensitive detection of proteins in plasma and serum samples. Antibody functionalized magnetic beads are used to capture the target of interest. Following capture, non-specifically bound proteins are washed off before PEA probes are added for detection of the bound proteins. Compared to hoogenous PEA, SP-PEA admits the use of larger sample volumes to increase available target molecules, higher concentration of detection reagents for more efficient formation of detection complexes and washes for removal of nonspecific background. We compared SP-PEA to solution phase PEA for the detection of cytokines: interlukin-6, interlukin-2, interlukin-4, interlukin-10 and Tumor Necrosis Factor-alpha, and we demonstrated an increased sensitivity by 15 to 60 fold in buffer and chicken serum. We further expanded SP-PEA to detect extracellular vesicles (EVs) through combinations of proteins on the surface of specific EV populations.

## Introduction

The complexity of the plasma proteome necessitates the demand for new analytical techniques to enable biomarker detection and discovery at the lower end of the abundance spectrum.^1^ In particular, early diagnosis and improved prognosis of disease can benefit from detection of lower levels of biomarker proteins, potentially unburdening healthcare systems and improving the quality of life for affected individuals. Proteins released from diseased tissue are present in minute amounts in the blood, usually as low as 10^10^ fold lower than the most abundant functional blood proteins such as Albumin.^2^ This disparity complicates detection of the blood biomarkers using immuno-assays and increasing cross reactivity during antibody detection which reduced the overall assay sensitivity. As such, higher sensitive tests can significantly expand opportunities for early diagnosis of diseases by detecting early signs of rising levels of plasma proteins indicative of disease in blood samples. Molecular protein detection assays with improved proofreading are needed to allow efficient detection of target molecules in a sample without troubling levels of nonspecific background as a means to augment specificity and sensitivity of detection. Improved protein detection techniques, with augmented sensitivity of detection, can greatly expand the scope for disease diagnosis by increasing the numbers of target molecules accessible for analysis, and potentially permitting detection at earlier stages of disease, where promising marker proteins may be present in even lower levels.

DNA-assisted proximity-based immunoassays have been developed for different proteomic applications, such as measuring protein expression^3^, posttranslational modifications^4,5^, visualization of protein in situ^6^, protein profiling^7,8^, detection of extracellular vesicles^9^, infectious diagnostics^10^ and western blotting^11^. Variant of proximity ligation assays have been developed where targets must be recognized by three antibodies, for enhanced detection and prostate derived microvesicles called prostasomes have been detected at elevated levels in plasma from prostate cancer patients using sets of five different antibodies.^12–14^ Multiplexed proximity extension assay (PEA) has become a standard research tool for analysis of multiple proteins require minimal sample volumes and attractive approach especially for high throughput detection of proteins in serum or plasma over broad dynamic range.^8,15,16^ In PEA, antibodies or other affinity reagents that are conjugated to oligonucleotides, such that target recognition by two or more such probes allow DNA sequence information on individual reagents to be brought together and combined into a single DNA strand via polymerization reactions. The resulting linear or circular DNA strands that form can subsequently be detected using efficient molecular genetic techniques. These proximity-based assays have proven a generally sensitive, specific, and efficient high-throughput technology for protein analysis. The method has been used and reported extensively for proteomic exploration of possible predictive and prognostic biomarkers in a wide range of diseases^17,18^, in protein profiling^19^ amongst other applications. With the quick present protocol and low volume significant advantages of the homogenous PEA assay, it nonetheless has some limitations in that there is no opportunity to remove unbound reagents. The sample volume may also contain ever lower amounts of target protein and cannot be easily expanded to increase the amount of target molecules.

Magnetic beads present significant opportunities as a support system that can be used and adapted in a variety of applications with improved assay performance.^20,21^ The use of magnetic particles in PEA could allow for controlled removal of unbound reagents and matrix components that could otherwise interfere with the assay. These include the possibility to use larger sample volumes with more target molecules, and to remove excess unbound and loosely bound reagents, allowing higher concentration of detection reagents for more efficient detection with little increase in background noise. Matrix effects from sample components that could otherwise interfere with the enzymatic reactions or with detection can also be minimized. This in turn allows us to analyze higher volumes of samples as we can reduce the possibility of background noise while increasing the likelihood of capturing a greater number of the target of interest as well the use of a capture antibody can decrease risks of cross-reactive detection.^22,23^

Here, we present a solid phase-based variant of the proximity extension assay (SP-PEA), which serves to improve the efficiency of detection to improve detection sensitivity and dynamic range of detection compared to homogenous PEA (Figure 1.). Target molecules are first captured from liquid samples via immobilized antibodies on magnetic beads. After washes pairs of PEA probes are added, followed by renewed washes to remove unbound reagents. Next, PEA probes that have bound in proximity allow hybridization of coupled oligos followed by extension of the 3’ end, the extension products are quantified via qPCR. We established assays for IL-6, IL-10, IL-2 and IL-4, demonstrating significantly increased sensitivity over PEA. The assay was also applied for detection of EVs from prostate cancer cell lines.

**Figure 1.**
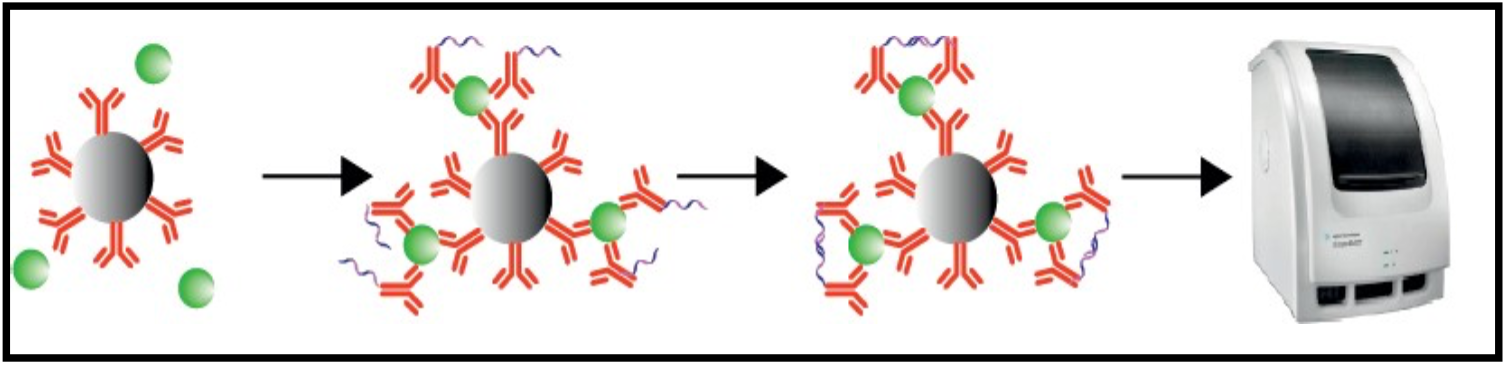
**A)** A Schematic representation of SP-PEA assay. I), antibodies are coupled to magnetic particles and unbound antibodies are removed by washes. II), the paramagnetic supports are then used to capture antigen from samples, remaining free antigens and other sample components are washed off. II), pairs of PEA probes (antibodies or other affinity reagents are conjugated to oligonucleotides) are then added to the reactions, incubated and free PEA probes are washed off. IV), only upon target recognition by the correct pair of detection reagent are DNA oligonucleotides brought into proximity to give rise to amplifiable and detectable extension products. The resulting DNA extension templates are then amplified and quantification by qPCR (5, 6).

## Materials and methods

### Antibodies, Antigens, Oligonucleotides and buffers

Unconjugated and biotinylated antibodies were purchased from R&D systems as shown in Supplementary Table 1. Dynal beads MyOne Streptavidin (10 mg/ml) were purchased from Invitrogen, USA (cat No. 65602). Buffers used include wash buffer (1xPBS with 0.1% BSA (New England Biolabs; NEB) and 0.05% Tween-20 (Sigma-Aldrich)), PEA buffer (1xPBS with 1% BSA, 100 μg/μL salmon sperm DNA (Invitrogen), 100 μg/mL goat serum (Sigma-Aldrich), and 0.05% Tween-20), and probe storage buffer (1xPBS, 0.1% BSA, 0.05% NaN3). All buffers were prepared in-house and their pHs were calibrated to 7.2. All enzymes, dUTPs and dNTPs were purchased from NEB. The PEA oligonucleotides conjugated to the antibodies are reported in Supplementary table 2. All oligonucleotide sequence was purchased from integrated DNA Technologies (IDT).

### Preparation of PEA Probes

PEA probes were prepared by coupling antibodies covalently to oligonucleotides. Antibodies were diluted in 1xPBS at 2 μg/μL and stored at −20°C until needed. Before conjugation, 4 mM DMSO was used to dissolve 25 mM dibenzylcyclooctyne NHS (DBCO-NHS) ester purchased from Jena Bioscience. The DBCO-NHS ester was 33.3-fold molar excess over antibodies. The 7K MWCO Zeba Spin columns purchased from Thermo Scientific were equilibrated before use according to the manufacturer’s instructions. Purified antibodies treated with DBCO-NHS ester were divided into two aliquots for preparation of the two PEA probes. 10 μg of DBCO-modified antibodies were mixed with a 2.5-fold molar excess of Forward PEA or Reverse PEA oligonucleotides and incubated overnight at 4°C. Conjugates were validated on 10% TBE urea denaturing gel and the PEA probes were diluted in PEA buffer before use.

### Solution Phase PEA

Recombinant antigen was diluted in PEA buffer. Pairs of PEA probes were mixed in the same buffer at a final concentration of 100 pM each, and 3 μl of probe mixture was added to 1 μl sample. After incubation at 37°C for 1 hour, 46 μl PEA extension/PCR mix (1xPCR buffer (Invitrogen), 0.1 μM each Forward PCR and Reverse PCR primers, 2.5 mM MgCl2, 0.2 μM TaqMan probe (or 0.5x sybr green I), 0.25 mM dNTPs (which have dUTPs), 0.02 U/μL Klenow exo-, 0.02 U/μL uracil-N-glycosylase, 1.5 U/μL Taq polymerase) was added. PCR was performed with the following cycles: 95°C for 2 min followed by 45 cycles of 95°C for 15 sec and 60°C for 1 min.

### Antibody Immobilization on Magnetic Beads

Dynal beads MyOne Streptavidin T1 beads were washed three times with 0,05% Tween 20 in 1xPBS pH7.4 (wash buffer), and biotinylated antibodies (final concentration 50 nM) were coupled to the beads by incubation at room temperature for 60 min. Unbound oligonucleotides were removed by two washes with wash buffer. A dilution series of the antigens was prepared in PEA buffer and 10% chicken serum (Invitrogen), and each dilution series included negative controls with no protein added to determine the assay background. In the assay, 45 μL of the sample was added to 5 μL of antibody coated beads and incubated for 60 minutes with rotation. The beads were then washed on twice with 2X volume of wash buffer on a magnetic rack.

### Solid-Phase PEA

A pair of PEA probes (final concentration of 500 pM each) were added to the beads and incubated for 1 hours at room temperature. PEA probes remaining in solution were washed as previously, followed by addition of 50 μL of extension mix (1X buffer 2 (NEB), 0.4 mM dNTPs, and 0.02 U/μL Klenow exo-) and incubation for 20 min at 37°C. Finally, a 2X PCR mix (2X PCR buffer, 0.2 μM each forward and reverse primer, 5mM MgCl2, 0.4 μM TaqMan probe (or 1X sybr green I), 1.5 U/μL platinum Taq polymerase and 0,4 nM dNTPs) was added to a final concentration 1X and PCR was performed with the following cycles: 95°C for 2 min followed by 45 cycles of 95°C for 15 sec and 60°C for 1 min.

### Data Analysis

The recorded Ct (cycle threshold) values for q-PCR data were further analyzed with Microsoft Excel and plotted as Ct values (y-axis) against protein concentration (x-axis). The data was analysed by linear regression to determine the LOD (limit of detection). The LOD was defined as the concentration of protein corresponding to CtLOD=CtN – (2 x SN), where CtN is the average Ct acquired for the background noise, and SN is the standard deviation of that value. 4-parametric linear regression was used to determine the LOD, LLOQ as well as extrapolating the sample concentrations using the Image J image processing software curve fitting function. The plots were generated on Microsoft Excel and R (http://www.R-project.org/).

## Results

### SP-PEA assay parameters optimization

In this study we set out to improve the sensitivity of the solution phase PEA by mainly focusing on three aspects; 1) Increasing the amount of detection reagents, 2) Washing off non-specific binding and 3) Increasing the sample volume. To investigate if increasing the detection reagents would result in increased signal without affecting the background. SP-PEA requires three antibodies that target different epitopes on the antigen of interest. We performed a comparison to see the viability of using wither monoclonal or polyclonal antibodies. Comparing monoclonal to polyclonal antibodies coupled to magnetic beads as capture reagents for IL-6 and IL-10, polyclonal antibodies performed better showing an improvement of 2 fold better sensitivity on both assays (Supplementary Figure 1.). To demonstrate the increased detection efficiency from using a higher concentration of detection reagents, we titrated for the optimum PEA probe concentration in the assay and Figure 2 shows the effects of variable PEA probe concentrations (500 pM, 1000 pM and 2000 pM) for the detection of IL-6. The increased signals obtained from using higher probe concentrations also came with elevated background levels, limiting the potential increase in sensitivity of the assay by increasing probe concentrations above 500 pM. As such we determined 500 pM to display the optimal signal to noise ratio which was then used for the rest of the experiments. Other conditions were also investigated for the most optimal solid phase PEA format including incubation time, temperature, rotation speed (data not shown).

**Figure 2.**
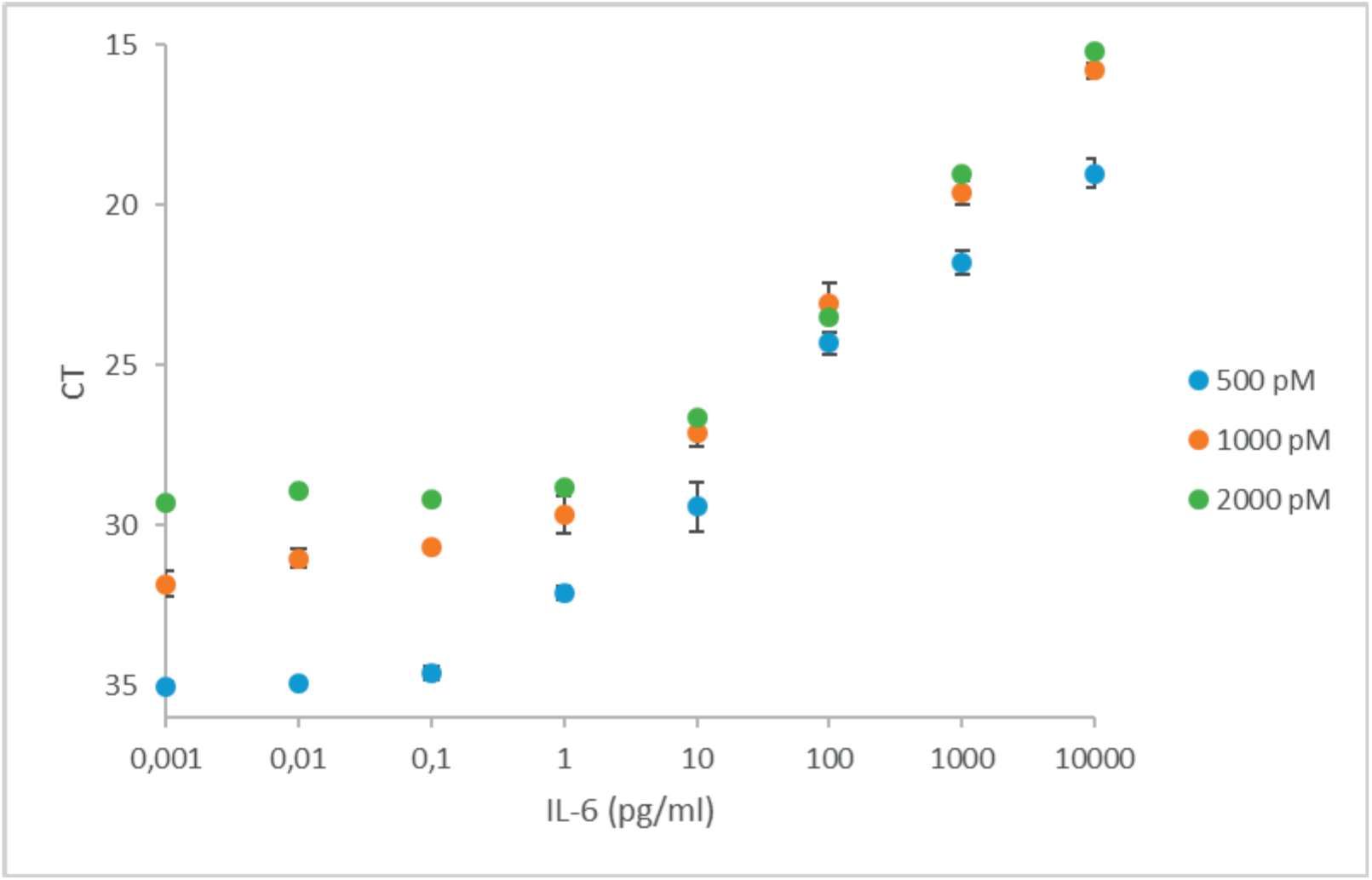
Optimization of probes concentration. Comparison of the effect of different probe concentrations 500 pM (blue), 1000 pM (orange), or 2000 pM (green) in buffer. The X-axis represents the antigen concentration in pg/mL and y-axis represents Ct from qPCR. All measurements were performed at least in triplicates. Error bars indicate SD.

### SP-PEA comparison to PEA

We compared the SP-PEA to solution phase PEA as reported by Lundberg *et al*., (3) for the detection of IL-6, IL-10, IL-4, IL-8 and TNF-alpha in buffer. The results are summarized in Fig. 3. The limit of detection (LOD) was calculated as described under data analysis in the Materials and Methods section and is represented in Table 1. When comparing the LOD for PEA to magnetic bead-based PEA, the lower limit of detection was improved by 1-2 orders of magnitude for the solid-phase variant, and the dynamic range was extended by a total 2-4 orders of magnitude. Furthermore, we observed a hook-effect for the homogenous PEA such that when target concentration increased beyond a certain point signals started decreasing again, while for SP-PEA a plateau signal was reached at higher antigen concentrations of IL-6 and IL-10. SP-PEA was capable of detecting lower concentrations for three of the investigated antigens (IL-6, IL-10 and TNF-alpha) compared with homogenous PEAs and with a broader dynamic range by up to 4 further orders of magnitude.

**Figure 3.**
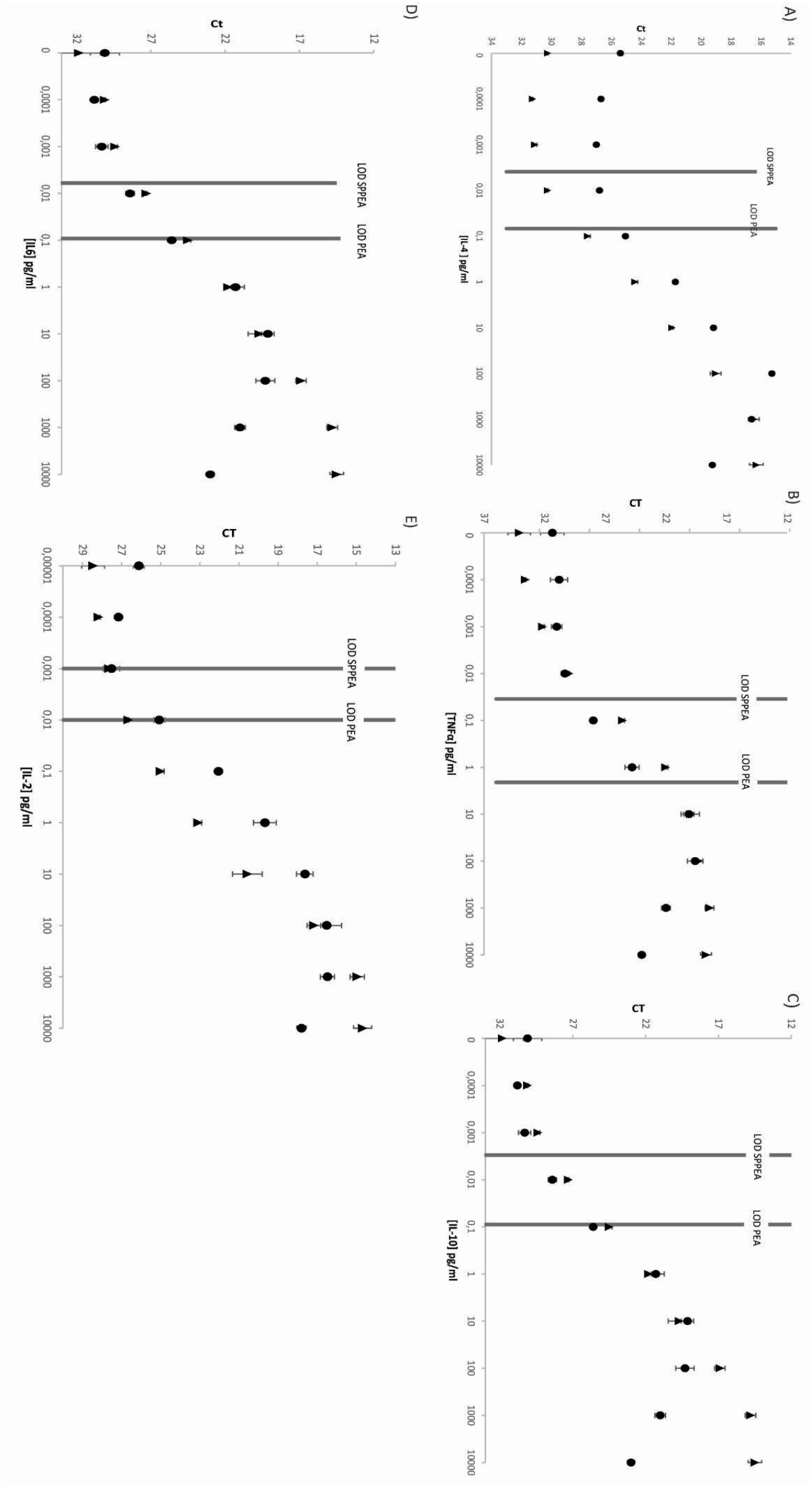
Assay performance comparison of SP-PEA (blue) with homogenous PEA (orange) for detection of IL-4, (B) TNFα, (C) IL-10, (D)IL-6 and (E) IL-2. A), comparison of detection of IL-6, B) IL-10 and C) TNF-alpha in assay buffer. X-axes represent the concentration in pg/mL and the y-axes represent Ct values.

**Table 1:**
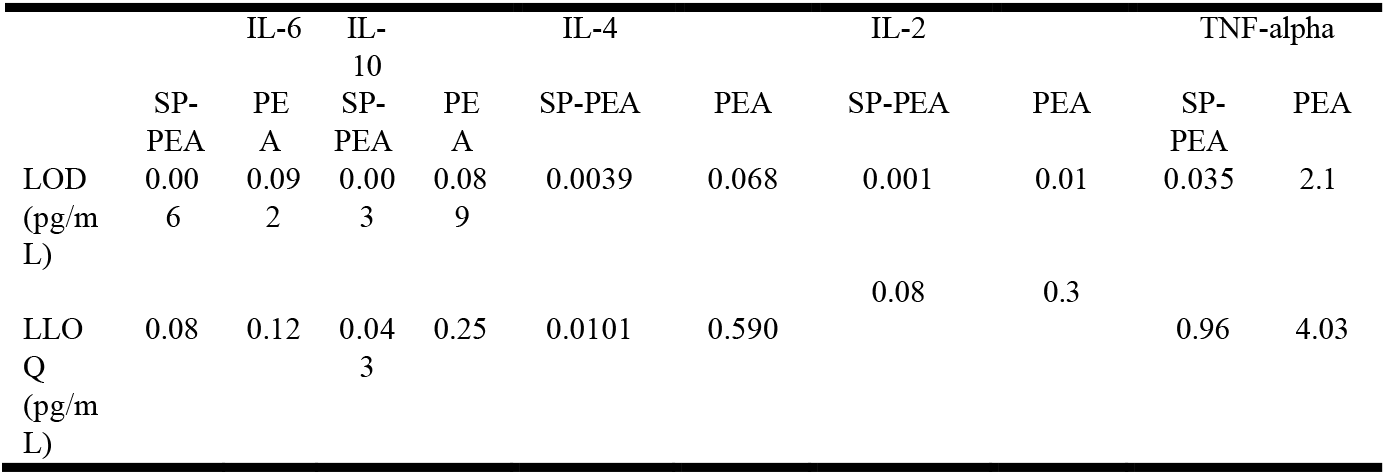
Comparison of LOD and LLOQ for SP-PEA in measurements of IL-6, IL-10 and TNF-alpha with homogenous PEA in buffer

|  | IL-6 |  | IL-10 |  | IL-4 |  | IL-2 |  | TNF-alpha |  |
| --- | --- | --- | --- | --- | --- | --- | --- | --- | --- | --- |
|  | SP-PEA | PEA | SP-PEA | PEA | SP-PEA | PEA | SP-PEA | PEA | SP-PEA | PEA |
| LOD (pg/mL) | 0.006 | 0.092 | 0.003 | 0.089 | 0.0039 | 0.068 | 0.001 | 0.01 | 0.035 | 2.1 |
| LLOQ (pg/mL) | 0.08 | 0.12 | 0.043 | 0.25 | 0.0101 | 0.590 | 0.08 | 0.3 | 0.96 | 4.03 |

### SP-PEA in complex samples

To investigate assay performance in more complex matrices, we investigated the detection of the two pro-inflammatory cytokines, IL-2, and IL-4, in dilution series prepared either in 10% chicken serum or in 50% chicken serum diluted in assay buffer using SP-PEA (Figure 4). We used chicken serum to represent the complex composition of human plasma or serum and the assay performance in serum was comparable to that in buffer. This illustrated that washing may have an effect on the removal of inhibitory substances that may be present in plasma samples. Assay characteristics in buffer, comparison of LOD, LLOQ and dynamic ranges for SP-PEA and homogenous PEA in measurements of IL-4 and IL-2 in buffer and in 10% chicken serum are summarized in Table 2. Assay performance is not affected in 10% chicken serum however at 50% we observed a reduction in assay sensitivity. The assay exhibited good inter-assay correlation and agreement with available clinical data as measured in a conventional ELISA (data not shown).

**Figure 4.**
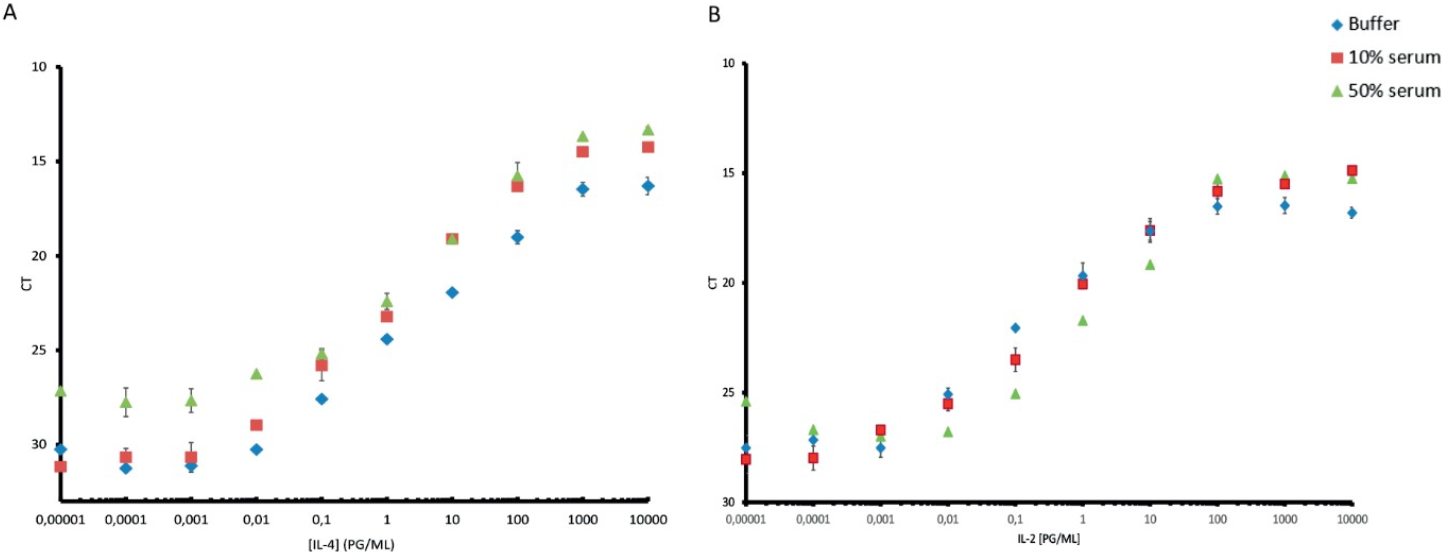
Comparison of SP-PEA performance in a complex matrix in 10% and 50% serum. Measurement of IL-4 and IL-2 in 50% chicken serum (green), 10% chicken serum (red) and in buffer (blue). X-axes represent the concentration in pg/mL and y-axes represent qPCR Ct values.

**Table 2:** Comparison of LOD, LLOQ and dynamic ranges for SP-PEA measurements of IL-2 and IL-4 in buffer and in 10% chicken serum.

|  | IL-4 | IL-2 |
| --- | --- | --- |
|  | SP-PEA | SP-PEA |
| Buffer LOD (pg/mL) | 0.0039 | 0.001 |
| 10% chicken serum LOD (pg/mL) | 0.09 | 0.2 |
| Buffer LLOQ (pg/mL) | 0.018 | 0.34 |
| 10% chicken serum LLOQ (pg/mL) | 1.28 | 0.8 |
| Dynamic range | 10<sup>6</sup> | 10<sup>5</sup> |

### SP-PEA detection of Prostate Cancer cell line derived extracellular vesicles

To demonstrate the suitability of SP-PEA triple recognition for extracellular vesicles (EVs) detection we detected serial dilutions of EV extracts from LNCaP human prostate carcinoma cells. We used combinations of three unique antibodies against markers known to be present on LNCaP derived EVs. Surface proteins commonly found on EVs, CD-9 and CD-81, were used as probes while PCa cell line specific marker, kallikrein as well as CD-38 were used as capture antibodies. The specificity of the EV detection was examined by using Calnexin as a negative protein marker. Significant detection was observed where CD-38 and kallikrein were used as capture antibodies while calnexin capture had only background signal. Detection limit of EVs using kallikrein as a capture was 80 ng/ml while using CD-38 as capture antibody had a detection limit of 20 ng/ml (Figure 5.).

**Figure 5.**
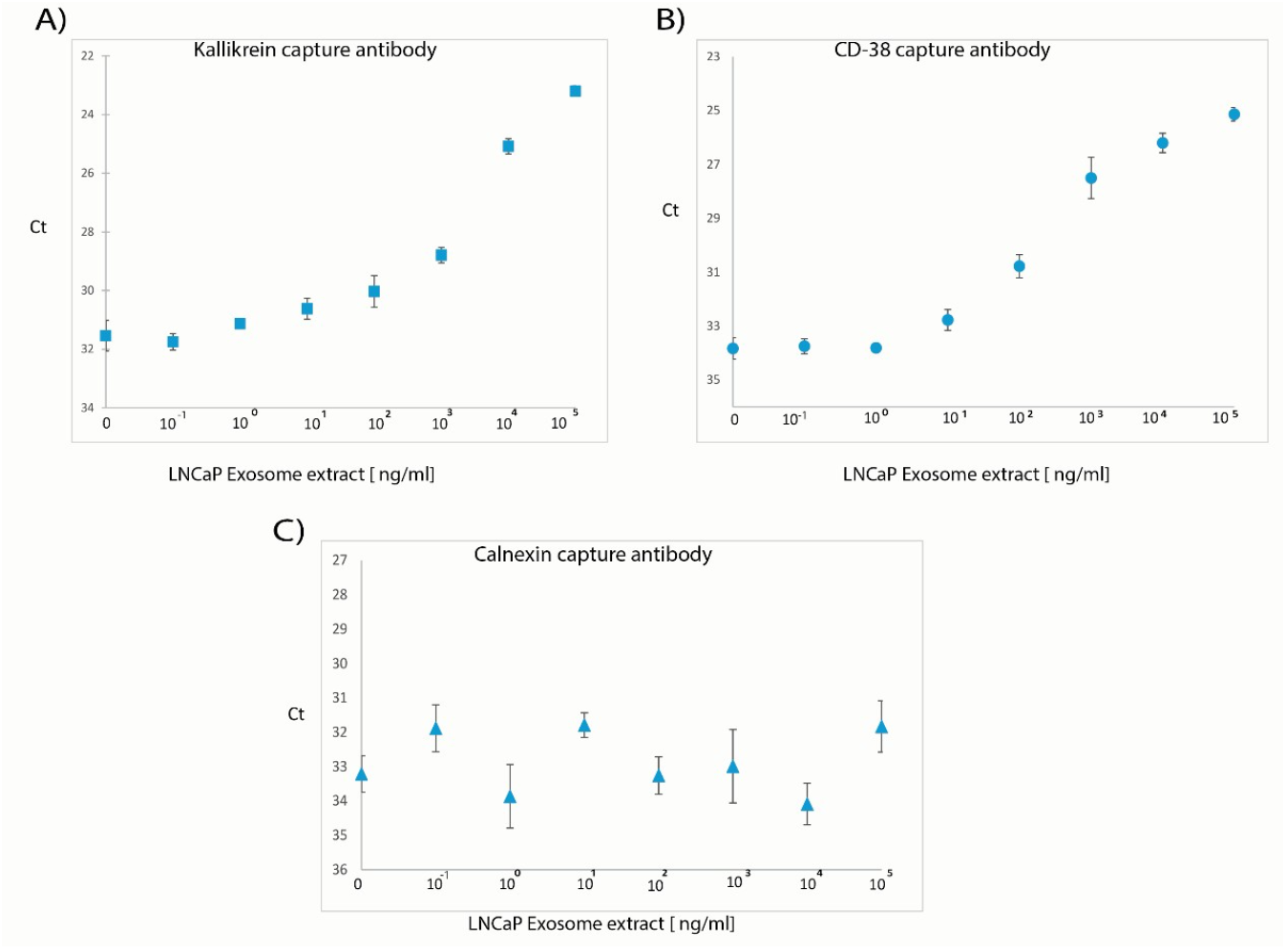
SP-PEA detection of serial dilution of EVs from LNCaP cell line with CD-9 and CD-81 antibodies using **A)** Kallekrein, **B)** CD-38 and **C)** Calnexin as capture antibodies. The y-axes show the qPCR Ct values while the x-axes show the concentration of LNCap derived EV extracts.

### Analysis of gastric cancer patient samples

Finally we validated the performance of SP-PEA with the detection of IL-4serum samples collected from gastric cancer patients (Figure 6.). Levels of IL-4 in serum samples of gastric cancer patients differed significantly from those of healthy controls (Wilcoxon signed-rank test, N gastric cancer patients = 24, N healthy controls = 25) when measured by both homogenous PEA (p-value < 0.0001) and SP-PEA (p-value < 0.0001). A considerable number of control samples were below the assay detection limit when analyzed by homogenous PEA as compared to analysis by SP-PEA.

**Figure 6.**
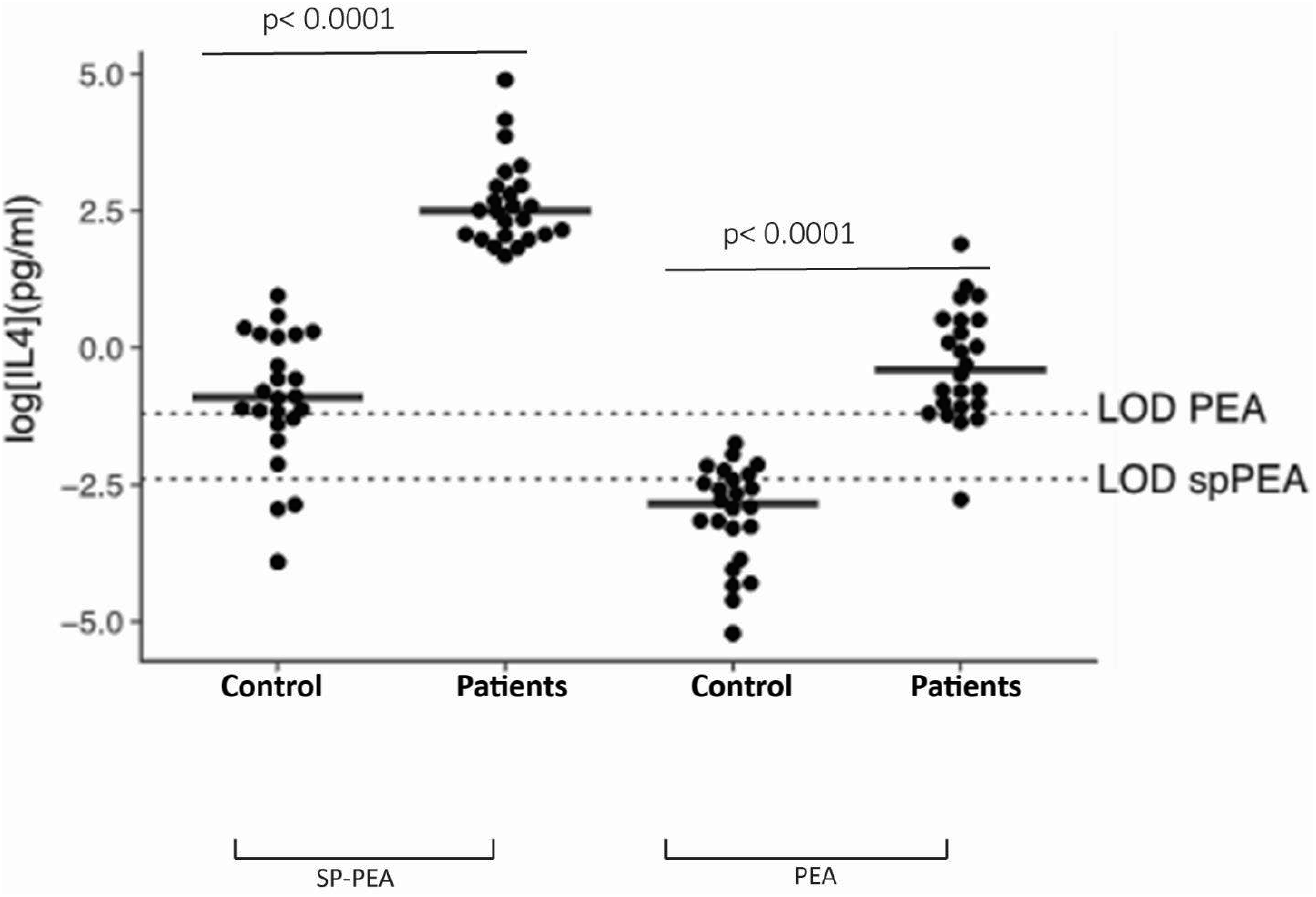
Comparison between PEA and SP-PEA for the detection of IL-4 protein levels in serum samples from gastric cancer patients (n =24) and healthy controls (n = 25). Each dot represents the mean of a triplicate. *P values* were calculated using a Wilcoxon signed-rank test. Respective limits of detection were calculated from standards included in the assays.

## Discussion

Homogeneous assays present significant advantages as no washes are required and small sample volumes can be used which is especially useful for precious biobank samples. Conversely, homogeneous assay has drawbacks in that unbound reagents cannot be removed, which contribute to background, limiting assay sensitivity and dynamic range. Thus, adding a separation step to remove all non-specifically bound reagents can allow the development of a more sensitive and specific immunoassay. Here, we report a magnetic bead-based PEA variant, which allows for increased assay volumes and removal of unbound reagents by washes and SP-PEA could detect lower protein concentrations compared to the standard homogenous PEA. However, the use of 50% serum as a diluent resulted in poor detection of target proteins (IL-6 and IL-4), perhaps due to aggregation of the particulate solid support. In 10% serum concentrations, we did not observe any significant difference in limits of detection of IL-6 compared to samples diluted in buffer. The addition of a solid phase for capturing targets protein from solution improved detection over solution PEA as shown in Table 1. This can be due to increased sample volumes with the corresponding increase in amounts of target proteins, or more efficient detection using the higher concentration of antibody-DNA conjugates, or some combination thereof. The requirement for target recognition also reduces the risks of crossreactivity for irrelevant target proteins through this additional proofreading step. Moreover, compared to ELISA only pairs of detection reagents can give rise to detection signals in SP-PEA, not individual detection reagents, thereby serving to reduce nonspecific background. Here, we showed that the dynamic range of SP-PEA was increased by three orders of magnitude for some analytes, and with lower detection limits reaching low femtomolar concentrations. Assay formats where solid supports are used require more user intervention, but have merits in that i) larger sample volumes may be used for increased sensitivity as needed, since washes are used to remove excess sample in place of the dilution used in homogeneous reactions. ii) matrix effects from sample components that could interfere with the enzymatic reactions or quench or nonspecifically add to the detected quality (fluorescence, absorbance, etc) can also be removed by washes. iii) It is also possible to use higher concentrations of detection reagents for more efficient detection, because surplus reagents are removed in the washes, thus preventing them from contributing to assay background. iv) Finally, the addition of detection reagent for capture brings the total number of recognition events required for proper detection of a protein to three. This has the effect of improving assay specificity over the more conventional assays depending on dual recognition, or in some formats using binding by single binders.

Cytokines as intercellular communication mediators are predicted to have important role in both inhibition and promotion of cancer growth.^24^ T-helper (Th)-1 type cytokines (IL-2 and interferon-γ) are pro-inflammatory and have shown antitumor activity and have been applied for clinical treatment of patients with melanoma and renal cell carcinoma.^25^ In contrast, Th-2 type cytokines (IL-4) are involved in inflammation-mediated carcinogenesis in human organs, including the gastrointestinal tract. As such, cytokine concentrations in patients have often been studied in cancer patients to determine correlation with disease outcomes. However, cytokine concentrations in gastric cancer patients are often below the limit of detection for conventional protein detection assays such as ELISA.^26^ Thus, the increased sensitivity in SP-PEA detection of low abundant proteins cytokines such as IL-4 can improve understanding of disease progression.

EVs are released by many different cell types, carrying with them biomolecules from the parent cells and thus can indicate the state of their tissue of origin.^27^ The membrane surface of such EVs are characterized by an abundance of integral membrane proteins such as CD-9, CD-63 and CD-81 while also containing tissue specific proteins that can be utilized to differentiate origin. Detection methods for EVs must therefore be able to sensitively detect EVs and distinguish different populations in biological fluids. Most current methods such as Flow cytometry require prior isolation of EVs or use only a single protein for identification.^28^ SP-PEA offers detection and identification of EVs through recognition by three unique markers, by substituting only the capture antibody the assay can be expanded to EVs from a wider array of cells or tissues.

To conclude, we have developed a magnetic bead-based PEA format, which has the potential to detect proteins at lower concentrations and over broader detection ranges compared to homogenous PEA. The assay also lends itself for detection of protein complexes as well as extracellular vesicles.

## Supporting Information

**Supplementary Table 1:**
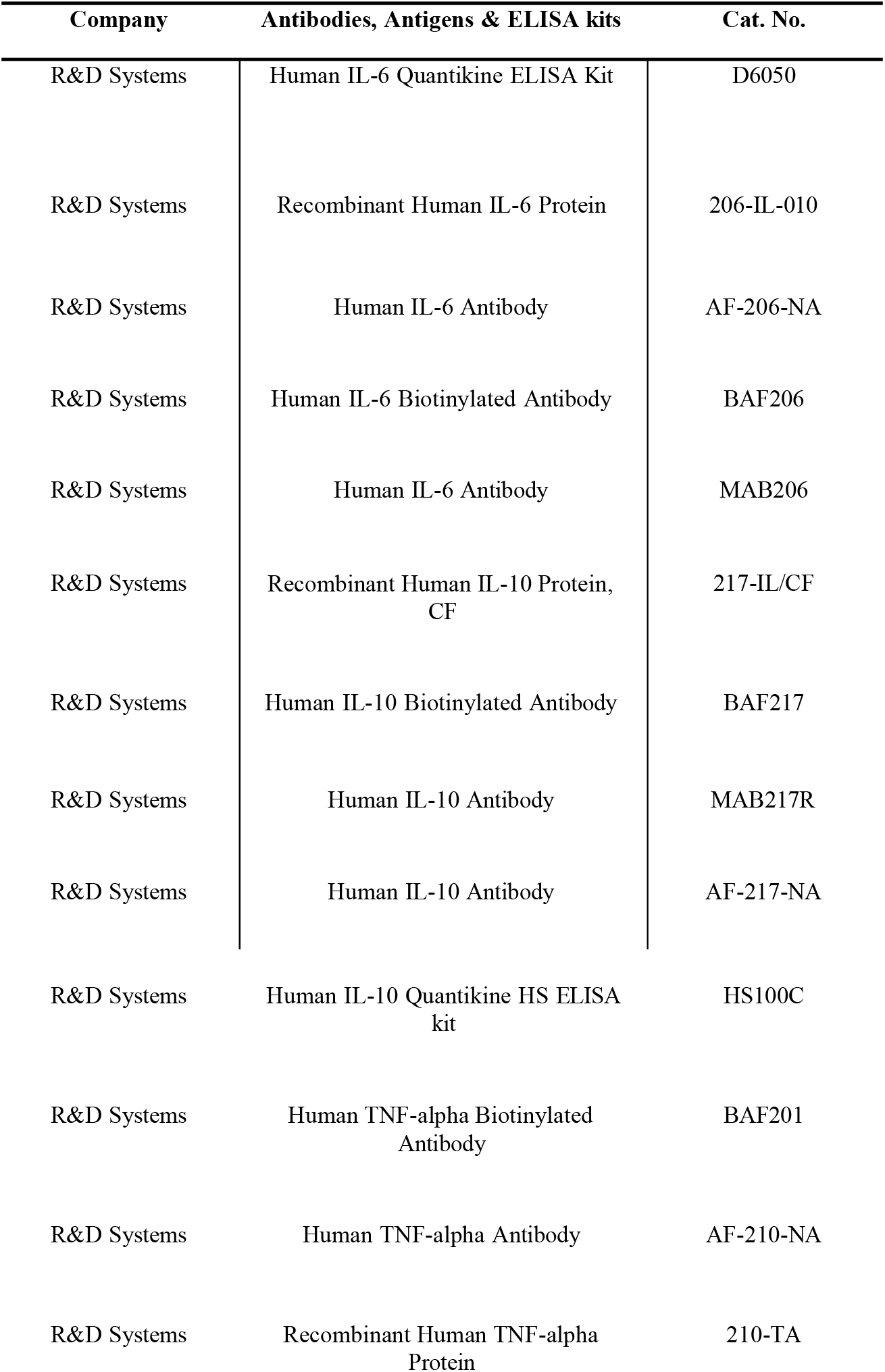
List of antibodies, antigens and ELISA kits

| Company | Antibodies, Antigens & ELISA kits | Cat. No. |
| --- | --- | --- |
| R&D Systems | Human IL-6 Quantikine ELISA Kit | D6050 |
| R&D Systems | Recombinant Human IL-6 Protein | 206-IL-010 |
| R&D Systems | Human IL-6 Antibody | AF-206-NA |
| R&D Systems | Human IL-6 Biotinylated Antibody | BAF206 |
| R&D Systems | Human IL-6 Antibody | MAB206 |
| R&D Systems | Recombinant Human IL-10 Protein,<br>CF | 217-IL/CF |
| R&D Systems | Human IL-10 Biotinylated Antibody | BAF217 |
| R&D Systems | Human IL-10 Antibody | MAB217R |
| R&D Systems | Human IL-10 Antibody | AF-217-NA |
| R&D Systems | Human IL-10 Quantikine HS ELISA<br>kit | HS100C |
| R&D Systems | Human TNF-alpha Biotinylated<br>Antibody | BAF201 |
| R&D Systems | Human TNF-alpha Antibody | AF-210-NA |
| R&D Systems | Recombinant Human TNF-alpha<br>Protein | 210-TA |

**Supplementary table 2:**
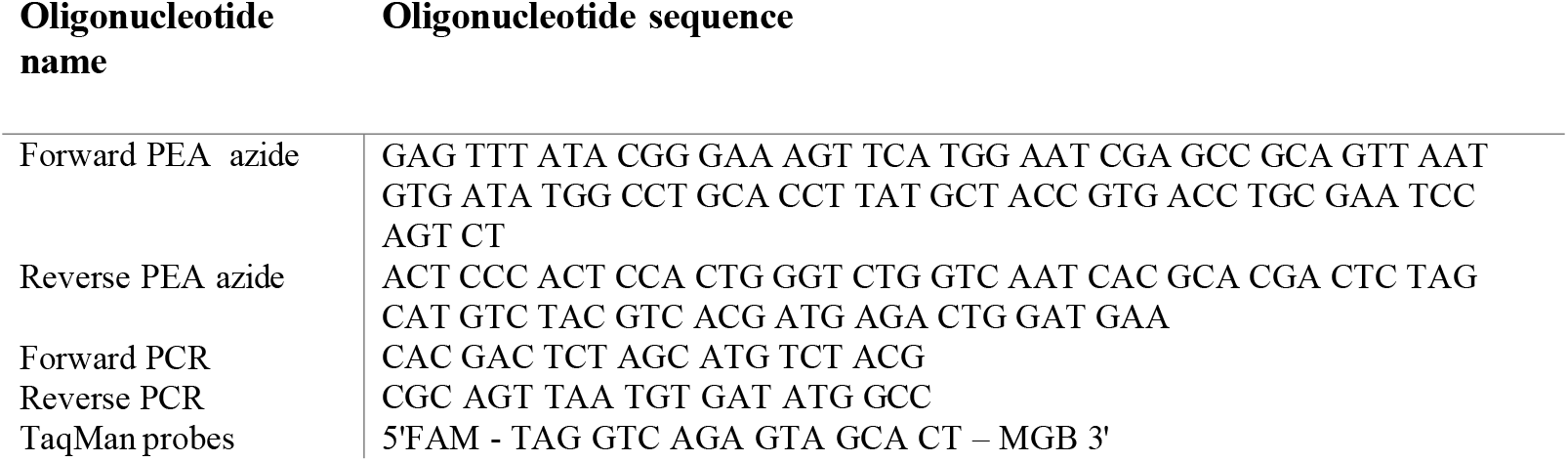
List of oligonucleotides

**Supplementary Figure 1.**
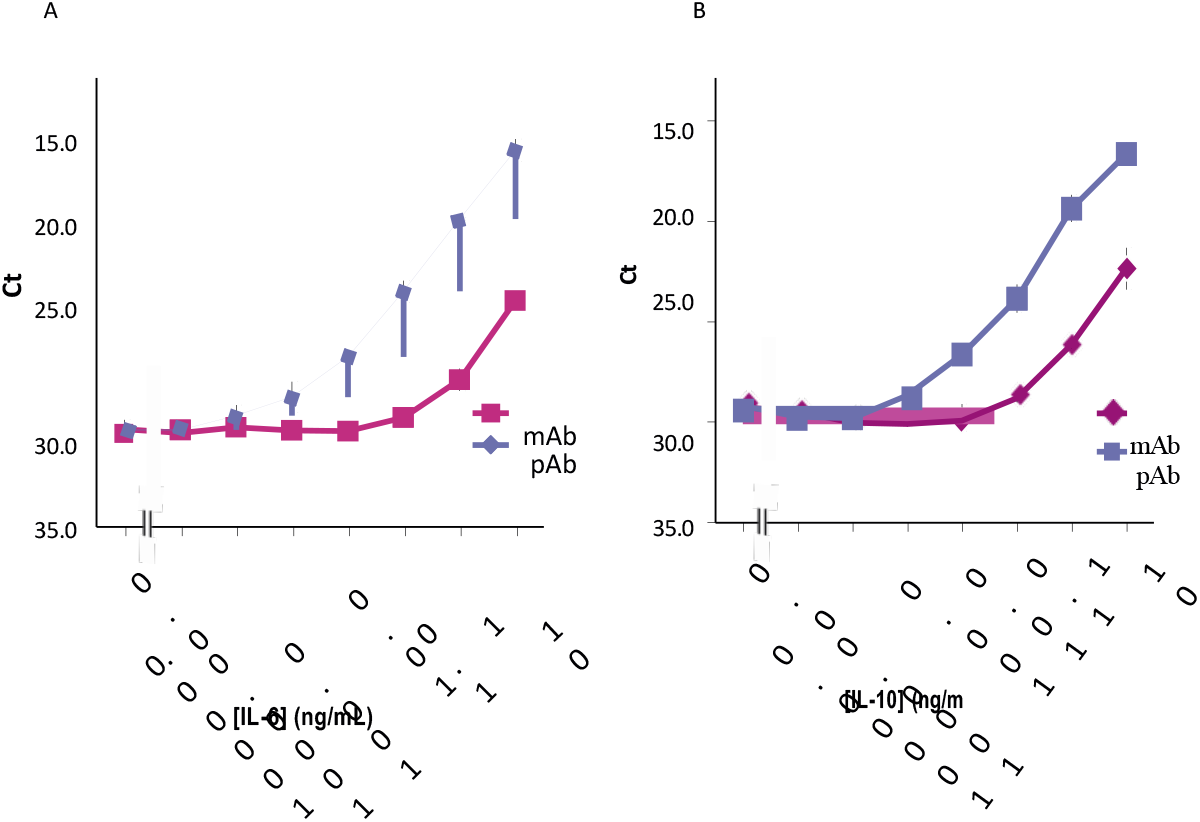
Comparison of the performance of monoclonal versus polyclonal antibodies as capture reagents coupled to paramagnetic beads for detection of (A) IL-6 and (B) IL-10. The X-axes represent the concentration in pg/mL and the y-axes represent Ct values.

